# Seed specific overexpression of a modified wheat *Or* gene leads to enhanced β-carotene in rice and wheat grains

**DOI:** 10.1101/2024.09.28.615562

**Authors:** Parul Sirohi, Ritika Vishnoi, Suchi Baliyan, Bidya Bhushan Gupta, Pratibha Demiwal, Hugo Germain, Debabrata Sircar, Harsh Chauhan

**Author notes:** Correspondence author: Harsh Chauhan, Department of Biosciences and Bioengineering, Indian Institute of Technology Roorkee, Roorkee, India.

## Abstract

Vitamin A deficiency is a major public health problem affecting up to 50% of the world’s population, as either wheat or rice, which are poor in many essential micronutrients such as vitamin A, are major staple food crops. Biofortification of cereal crops with β-carotene (provitamin A) through genetic engineering is a potential solution to overcome vitamin A deficiency. The Orange (Or) protein is involved in the regulation of carotenoid accumulation and previous studies demonstrated high carotenoid accumulation due to a single-nucleotide polymorphism (SNP) in the CDS leading to substitution of Arg to His in the OR protein results in carotenoid accumulation. In the present study, we showed that this substitution of a single amino acid at position 110 (Arg to His) of wild-type wheat *TaOr* (referred to as *TaOr^His110^*) increased β-carotene accumulation in transgenic wheat and rice plants overexpressing *TaOr^His110^* under the control of the seed-specific promoter *Glu-1D1*. HPLC analysis revealed increase in β-carotene content in rice grain up to 8-fold in case of TP309 (japonica) cultivar, 13-fold in case of IET10364 (indica) cultivar and 7-fold in wheat cv. CPAN1676. Additionally, most of the carotenoid biosynthetic pathway genes were found to be upregulated in *TaOr^His110^* overexpressing seeds of TP309 and IET10364, which positively correlated with maximum increase in β-carotene content.

## Introduction

The global population is estimated to exceed the milestone of 10 billion by the year 2050, and with increasing population come empty stomachs to feed (Sullivan J.N., 2023). As of now, there are roughly 2 billion people, suffering from hidden hunger, which is caused by not having a balanced diet providing all essential nutrients and not just carbohydrates (Muthayya et al., 2013; Gillespie et al., 2016). Though micronutrient deficiency can happen to anyone, children (43%) and pregnant women (38%) are more prone because of the increased nutritional demand in pregnancy and growth. The number of preschool-age children who suffer from malnutrition is around 20 million, largely from South-East Asia and Africa (Uauy et al., 2012), and about five million children under the age of five are dying every year because of malnutrition (FAO report, 2013). Among adults, micronutrient deficiency is a major cause of early death, mental retardation and blindness for almost one-third of the developing world’s population. The predicted economic loss because of malnutrition and micronutrient deficiencies may cost the world 2% to 3% of GDP, or US$1.4 trillion to US$2.1 trillion each year (FAO, 2013).

Micronutrients are essential for the normal growth of the human body as they are required for both physical as well as mental development (Godswill et al., 2020). They maintain essential functions and primary metabolic processes by functioning as cofactors of different enzymes and thereby. Since, the body cannot generate the required micronutrients, it is necessary to fulfill this demand through the diet. According to the United Nations World Health Organization, for the developing countries, famines are not as challenging as the poor nutrition lacking in micronutrients, resulting in hidden hunger. Important micronutrients required for proper growth and development are vitamins (A, B, and C), iodine, iron, and zinc. Vitamin A is an absolute requirement in human diet and Vitamin A Deficiency (VAD), the reduction in serum vitamin A levels caused by low dietary consumption of vitamin A or its precursors, causes, leads to decreased immunity and higher occurrence of infectious diseases, such as measles, intestine and lung infections, and impaired metabolism and tissue development (World Health Organization, 2009).

Crop biofortification refers to the production of a crop with enhanced nutritional value, this can be accomplished through conventional breeding methods or through genetic manipulation. In contrast food supplementation, biofortification emerges as better and more economically sustainable method, as once a crop is developed no further cost is incurred to produce micronutrient rich food. For staple crops biofortification, one of the most prevailing methods is targeted breeding, which involves the crossing of existing micronutrient rich varieties.

However, the complete non-availability of iron or provitamin A in staple cereals such as rice makes it impossible to enrich rice for these micronutrients using conventional breeding. Additionally, low heritability and lack of genetic diversity for micronutrients biosynthesis pathways is another challenge encountered in breeding. In these scenarios, transgenic technology comes to the rescue as it breaks these barriers and suitable genes can be introduced. The most popular example is the beta-carotene biofortified rice popularly called ‘Golden Rice’ (Ye et al., 2000; Paine et al., 2005). Biofortified staple food crops have also been developed for many micronutrients such as iron, zinc, iodine, vitamin B, C and E (Malik and Maqbool., 2020).

There are three strategies used for transgenic production of pro-vitamin A carotenoid in staple food crops. The first strategy, often known as the ‘push strategy,’ overexpresses genes encoding for the rate-limiting stages in the pro-vitamin A biosynthesis pathway. Golden Rice, which has multiple genes introduced, is an example of this strategy (Ye et al., 2000). Secondly, the blocked strategy silences a biosynthetic step that occurs immediately after the metabolic component whose level has to be raised. For example, Silenced lycopene-cyclases resulted in higher lycopene accumulation in tomatoes (Diretto et al., 2006). The third and most recent strategy, known as sink engineering, involves managing carotenoid storage structures to ensure that carotenoids are stored in a stable manner (Diretto et al., 2010; Li et al., 2012). One example of sink engineering is the use of *Or* gene, initially identified from an Orange cauliflower mutant (Zhou et al., 2008).

A novel *Or* gene isolated from an orange cauliflower mutant with high β-carotene has been proposed for provitamin A biofortification in both rice and wheat endosperm. Expression of *Or* gene mutants containing an insertion of a copia-like LTR retrotransponson or of wild-type *OR* with a single amino acid change have been demonstrated to significantly increase carotenoid accumulation in usually low-pigmented tissues (Zhou et al., 2008; Yuan et al., 2015). Lopez et al. (2008) demonstrated that by manipulating sink formation, the *Or* gene can increase carotenoid levels in transgenic potato tubers. *OR* is a highly conserved plant gene, encoding for a DnaJ Cys-rich zinc finger domain containing protein (Lu et al., 2006). Interestingly, the functional genomic investigation of a melon *OR* gene (*CmOR*) showed that a single nucleotide polymorphism resulting in an Arg to His change at position 108 was found to be contributing the orange phenotype of the fruits (Tzuri et al., 2015).

In the present study, we have employed genetic engineering editing approaches involving the wheat *Or* homolog to try to increase provitamin A content in both rice and wheat. We employed site directed mutagenesis to introduce the critical His substitution in the Or protein and used an endosperm specific promoter to target the modified *Or* gene expression in rice and wheat endosperm.

## Materials and methods

### Materials

#### Plant Material

The experiments were conducted to increase β-carotene concentration in rice and wheat endosperm through overexpression of mutated wheat *Or* (*TaOr*) gene (Genebank accession no. AK457010.1, cDNA accession no. Traes_6AL_9CEDE1FC0.1) in rice and wheat, followed by biochemical analysis of transgenic seeds. For plant material wild-type plants were grown in a greenhouse facility and under open conditions in potted soil during growing season at the Department of Biosciences and Bioengineering, IIT Roorkee.

#### Cloning of seed-specific promoter *Glu-1D1*

DNA isolation from leaf tissue was done by using the CTAB method as described by Yu et al., (2017). Quality of isolated DNA was checked on 0.8% agarose gel prepared in TBE buffer and DNA quantification was done using a Nanodrop (Thermo Scientific, USA). For the amplification of the *ProGlu-1D1* promoter (Gene bank accession no. AJ301618.1, Lamacchia et al., 2001), a PCR was performed using wheat genomic DNA as the template, *SwaI-Glu1D1*-F and *SpeI-Glu1D1*-R primers (Supplementary Table S1) and Phusion^TM^ DNA polymerase enzyme (Thermofisher Scientific, USA). The product was cloned in *pJET1.2*/blunt PCR cloning vector to make *pJET1.2-ProGlu-1D1*, which was sequenced before further experiments.

#### Cloning of *TaOr* cDNA, site directed mutagenesis for *TaOrHis^110^* and construct assembly in binary vector *pANIC6B*

RNA isolation from wheat leaf tissue was done by using Trizol reagent as per manufacturer’s protocol (Invitrogen, Thermofisher USA). 2µg RNA and *TaOR-F, TaOR-R* primers were used for one-step cDNA synthesis and PCR amplification of the 984 bp ORF of *TaOR* gene using one-step SuperscriptII RT-PCR system kit (Invitrogen, Life Technologies, Rockville, MD, USA) according to manufacturer’s instructions. The resulting PCR product was then cloned in the *pJET1.2*/blunt PCR cloning vector to generate *pJET:TaOR*, which was sequenced. Site-directed mutagenesis of wheat *OR* gene was done for the single amino acid substitution (Arginine to Histidine) at 110^th^ position by point mutation of CGC (Arginine) to CAT (Histidine). PCR was done using *pJET:TaOR* plasmid as template, *MutTaOr-F* and *MutTaOr-R* primers (Supplementary Table S1) and Phusion^TM^ DNA polymerase enzyme (Thermofisher Scientific, USA). After PCR amplification, the entire reaction mix was digested with Dpn1 (to degrade the parental strands). Digested product was used for transformation of DH5α competent cells. The site-directed mutagenesis and integrity of *TaOR* to *TaOr^His110^*was confirmed by plasmid sequencing.

Subsequently, subcloning of *Spe*1*-HindIII-TaOr^His110^*under seed-specific promoter *SwaI-Spe*1*Glu1D1* in binary vector *p6oAct-UbiZm-LH* was done by using specified restriction enzymes, to create binary vector *p6OAct-proGlu1D1-TaOr^His110^*. First, the Pro*Glu1D1* was taken out from *pJET1.2* using *Swa*1*-Spe*1 and replaced with *ZmUbi* promoter in *p6oAct-UbiZm-LH*. Second, *SpeI-HindIII-TaOr^His110^* was cloned downstream of promoter *Glu1D1* to generate *p6OAct-proGlu1D1-TaOr^His110^* binary plasmid.

Finally, the fragment *SpeIProGlu1D1:HindIIITaOr^His110^*containing entire cassette with *Promoter Glu1D1* and *TaOr^His110^ ORF* was excised by Swa1-EcoRV restriction digestion and cloned in the modified binary vector *pANIC6B* for *Agrobacterium* mediated transformation of both wheat and rice. The complete cassette cloned in *pANIC6B* is represented in Supplementary Fig. 1. Cloning and plasmid integrity was confirmed by sequencing.

#### Construct assembly of *TaOr^His110^* in *pB7FWG2* for subcellular localization

For subcellular localization, first the entry clone was created by cloning *TaOr^His110^* (without stop codon) with flanking attB sites into *pDONR221,* by using *TaOr-GFP-F* and *TaOr-GFP-R* primers (Supplementary table S1) and Gateway™ BP Clonase™ II Enzyme mix (Invitrogen) as per manufacturer’s instructions. Subsequently, sub-cloning of *TaOr^His110^*from *pDONR221* to *pB7FWG2* (destination vector) was done by performing the LR reaction using LR Clonase™ enzyme mix (Invitrogen) as per manufacturer’s instructions to generate the construct *pB7FWG2:35S:TaOr^His110^:*GFP. The cassette was confirmed by sequencing.

#### *Agrobacterium* infiltration of *Nicotiana benthamiana* and subcellular localization

For transformation, *A. tumefaciens GV3101* harboring the binary vector *pB7FWG2:35S-TaOr-GFP* was grown in LB-broth media containing 50 mg/L of spectinomycin, 30 mg/L gentamycin, 30 mg/L of rifampicin to the stationary phase at 28 °C. After centrifugation, *A. tumefaciens GV3101* was resuspended in the infiltration medium (10 mM MgCl_2_, 100 μM acetosyringone) to a final OD_600_ of 0.5. The suspension was infiltrated with a 1-mL needleless syringe into the abaxial surface of 4-week-old leaves of *N. benthamiana*. The *TaOrHis^110^ GFP* fusion localization was analyzed 24-48 hours after infiltration by confocal laser scanning microscope (TCS SP8, Leica). GFP was excited at 488 nm and the fluorescence detected between 498-525 nm.

### Genetic transformation of rice cultivars TP309 (japonica) and IET10364 (indica) and Wheat cultivar CPAN1676 using *Agrobacterium*-mediated transformation method

Rice transformation was done by *Agrobacterium*-mediated method as described by Toki *et al*., (2006) by using mature seeds of TP309 and IET10364 as explants. For wheat transformation, we followed the *Agrobacterium*-mediated method described by Hayta et al., (2019) using immature embryos as explants.

### GUS histochemical assay

Verification of transgenic plants was done by using GUS histochemical assay. For histochemical analysis small pieces of leaf tissue were inoculated in 1.0 ml GUS staining buffer (100 mM sodium phosphate buffer, pH-7.0, 100 mM potassium ferrocyanide, 100 mM potassium ferricyanide, 0.1% triton-x-100) containing 1 mM X-Gluc (5-Bromo-4-chloro-3-indolyl-Beta-D-glucuronide)) (Jefferson et al., 1987), vacuum infiltrated for 20 min and incubated at 37 ◦C for overnight.

### Molecular analysis of transgenic plants

To examine the integrity of *TaOr ^His110^* gene in rice and wheat genome, genomic DNA from the leaf tissue of transgenic plants was isolated using the Qiagen DNeasy Plant Mini Kit according to manufacturer’s protocol. A PCR amplification reaction was performed for each candidate plant using gene specific primers of *TaOR* and the selectable marker gene encoding *HptII* (listed in Supplementary Table S1). The amplified PCR products were run on 1% agarose gel and visualized on a gel documentation system. PCR positive plants were further analyzed for overexpression. Total RNA was isolated from leaf tissues and immature seeds of all the T0 plants. RNA isolation from two-week-old seedlings was done using the SV total RNA isolation system (Promega, USA) according to manufacturer’s instruction, including an on-column DNase treatment. RNA from developing seeds was isolated by using methods described by Singh et al. (2003) for carbohydrate rich seeds followed by RNA purification with the SV total RNA isolation system (Promega, USA).

First strand cDNA was then synthesized from 1 µg of total RNA using the Go-Script reverse transcription system (Promega, USA) as per manufacturer’s instructions. qPCR analysis was performed using *Ubiquitin5* (Jain et al., 2006) and Actin (Choi et al., 1991) as internal standard in rice and wheat respectively. The relative fold change in expression was calculated according to 2^^-ΔΔCt^ method. To ensure a stable expression, a similar analysis of T2 plants generated from confirmed T1 transgenic lines was also performed.

### Carotenoid extraction from rice and wheat grains

0.3 g dried seeds of both rice and wheat were ground in mortar and pestle and suspended in 2.5 ml of n-hexane:dichloromethane (1:1) followed by centrifugation at 9000 rpm, 5°C for 10 minutes. After centrifugation, the organic phase was collected and 500 µl of methanolic KOH (400 g/l) was added to each sample. Samples were kept at 50°C for 1 h. 0.5 volume of H_2_O and 0.5 volume of sodium sulfate (100 g/l) were added to the solution, mixed for 5 min, and centrifuged. The organic phase was collected and dried by centrifugal evaporation at 30°C. The pellet was resuspended in 250 µl acetonitrile and analyzed using HPLC-PDA.

### HPLC chromatographic conditions and analysis of β-carotene level

The chromatographic analysis of rice and wheat samples was performed on a Waters HPLC system. A X-bridge C18 (150 × 4.6 mm, particles 5 *μ*m) chromatographic column was used. The PDA detector was set to a wavelength of 499 nm. Gradient elution with a flow rate of 1 ml•min^−1^ was used. The column oven was conditioned at 25°C, the injection volume was 20 *μ*l and the analysis time was 23 minutes. With this method, β-carotene detected at 18.3 min.

The system consisting of a 1525 binary pump and 2998 photodiode array detector (PDA) (Milford, MA, USA). The PDA detector was set at 198-700 nm for recording chromatograms. Chromatographic separation was performed on the C18 reversed-phase column SunFire (150 x 4.6 mm, 3.5µm) of Waters (Milford, MA, USA). The isocratic mobile phase consisted of a ratio of 50:50:2 acetonitrile, methanol, and 2,4-dichloromethane were used to separate the analytes at a flow rate of 0.7 ml•min^−1^. The sample injection volume was 20 µL. Data acquisition and analyses were performed by Empower 3 Software from Waters. The total analysis time per sample was 35 min. HPLC chromatograms were detected using a photodiode array detector at 450 nm. The *β*-carotene present in the sample was identified by matching its retention time and UV-spectrum with those of authentic *β*-carotene standards under the same HPLC run conditions. The *β*-carotene was quantified by comparing the peak area of the standard β-carotene of known concentration.

### LC-MS analyses of *β*-carotene

Mass spectrum was acquired through ACQ-QDA Instrument Setup (Waters, Milford, MA, USA). The following run conditions were used: Positive capillary voltage (0.87 kV), positive cone voltage (15 kV), sampling rate (3.3 points/sec), start mass (115 Da), end mass (1100 Da).

## Results

### Cloning and site-directed mutagenesis (SDM) of the *TaOr* gene

In the present investigation we performed site-directed mutagenesis of *TaOr* gene for single amino acid substitution of Arginine (CGC) to Histidine (CAT) at 110^th^ position. The nucleotides 329 and 330 of *TaOR* were changed from GC to AT by SDM, resulting in the codon CGC to CAT alteration and changing Arg to His in *TaOR*.

### *TaOr* is a chloroplast-localized DnaJ-like cysteine-rich domain-containing protein

*TaOR* gene from wheat encodes a predicted protein of 328 aa, which is a DnaJ-like cysteine-rich domain-containing protein along with a transmembrane domain (Fig. 1A). Therefore, to assess the subcellular localization of *TaOr^His110^*, a C-terminal GFP fused construct with *TaOr^His110^* was prepared and transiently expressed in *N. benthamiana* leaves using agroinfiltration. Epidermal cells of infiltrated leaves were examined by confocal laser scanning microscopy. The results showed that GFP fluorescence of GFP: *TaOr^His110^* fusion protein was found to be localized in chloroplasts (Fig. 1B).

**Fig. 1.**
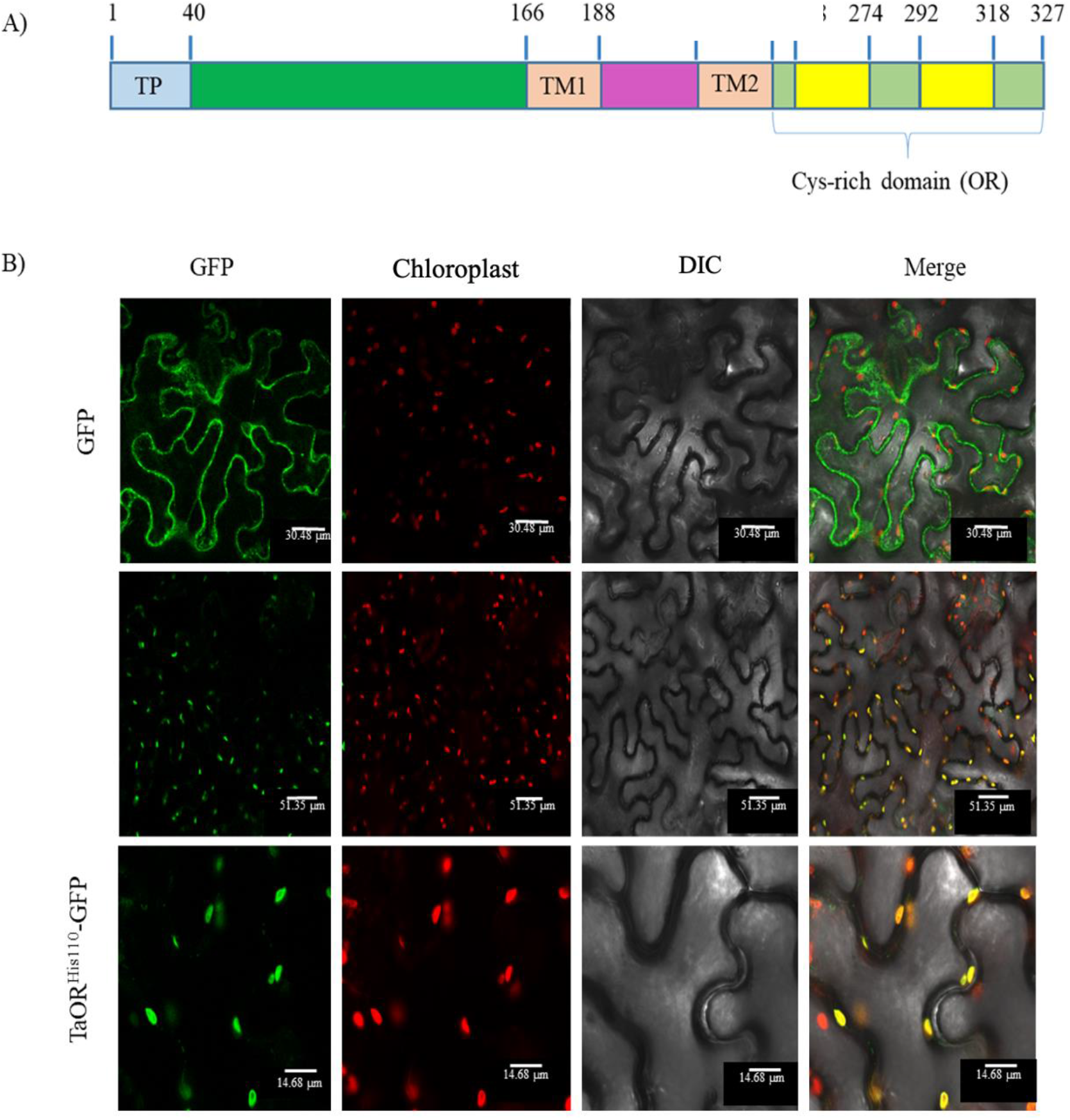
*TaOr* accumulates in the chloroplasts (A) Modular structure of *TaOr* (B) C-terminal GFP fused construct *TaOr^His110^* GFP: localization in *N. benthamiana* cells, upper panel GFP control, central panel MuTaOr at low magnification, lower panel MuTaOr at higher magnification

### Generation and verification of rice transgenic lines

We developed 60 transgenic lines in rice cv. TP309 and 25 in IET10364 using *A. tumefaciens* strain EHA105 carrying binary vector *pANIC6B:Glu1D1-TaOr^His110^.* The putative transgenic T0 plants were removed from the non-transgenic plant using HPT, gene specific PCR (Supplementary Fig. S3A and B) and GUS histochemical assay (Supplementary Fig. S2)

### *TaOr^His110^* is expressed in developing rice seeds

qPCR analysis was performed to analyze the expression pattern of transgene in T1 and T2 seeds. The relative expression of *TaOr^His110^*gene showed expression level in transgenic lines while the wild type didn’t have any expression. (Fig 2A). The maximum 1732- and 650-fold high expression of *TaOr^His110^*gene was observed in PS38 (Fig. 2A) line of TP309 and PS10 (Fig. 2B) line of IET10364 respectively. Further, in T2 seeds 20 days after anthesis higher expression is obtained as compared to wild type control (Fig. 2C and Fig. 2D).

**Fig. 2:**
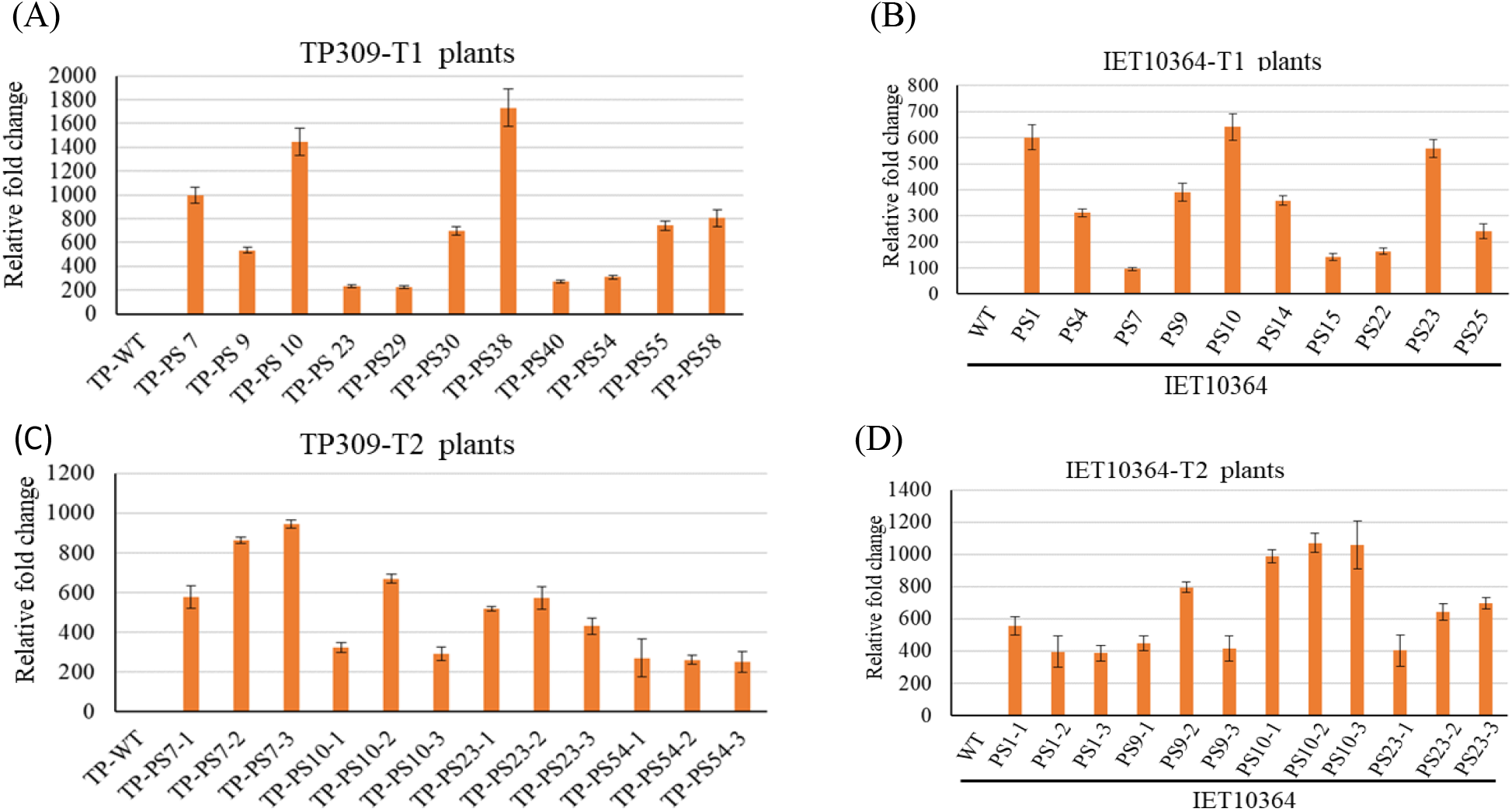
Relative expression of *TaOr^His110^*in developing seeds of *Glu1D1TaOr^His110^* transgenic japonica TP309 plants (A-C) Expression analysis in T1 generation with WT control. (B-D) Expression analysis in T2 generation.

### Estimation of β-carotene concentration in the grains of TP309-*Glu1D1TaOr^His110^* and IET10364-*Glu1D1TaOr^His110^* rice lines grains

A clear distinction in color between transgenic seeds and wild type seeds could be observed (Fig. 3). Further estimation of β-carotene concentration of four high expressing lines of IET10364-*Glu1D1TaOr^His110^* and five lines of TP309-*Glu1D1TaOr^His110^* transgenic (T2) rice grains were performed through HPLC. The sample chromatograms for β-carotene standard, transgenic seed and wild type seed are shown in Fig. 4. The increase in β-carotene concentration of transgenic grains of IET10364 ranged from 4.87 µg/g (IET10364-PS4) to 13.54 µg/g (IET10364-PS23). In IET10364 WT seeds, the β-carotene was either absent or below the detection limit. Similarly, in TP309 WT seeds the β-carotene was either absent or below the detection limit whereas β-carotene concentration of transgenic grains of TP309 ranged from 2.82 µg/g (TP-PS55) to 8.20 µg/g (TP-PS7). (Table 1).

**Fig. 3:**
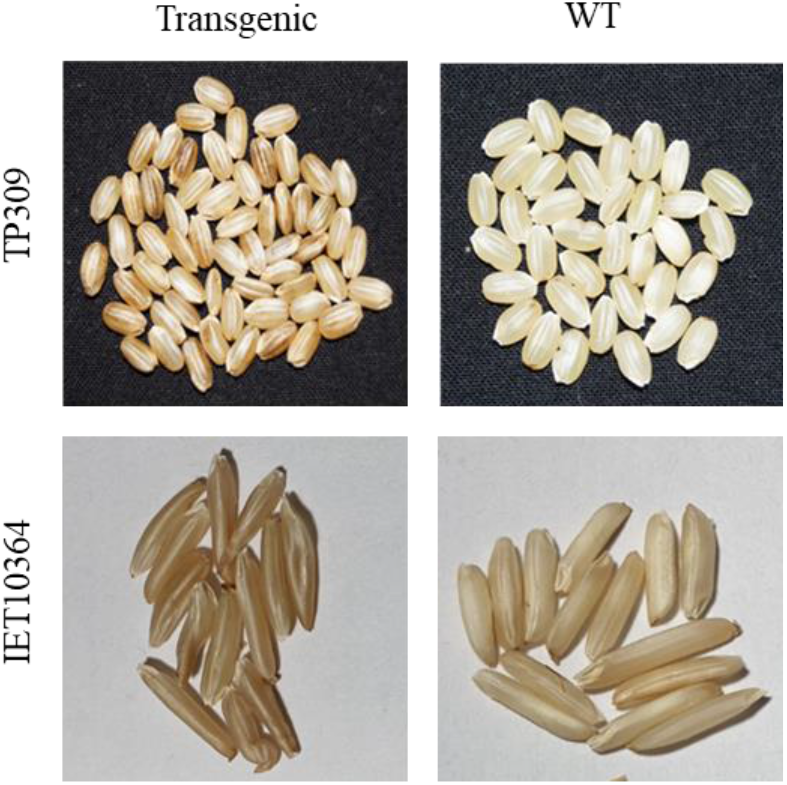
Comparison of transgenic seeds of TP309-*GluTaOr^His110^* and IET10364-*GluTaOr^His110^*with their respective wild type seeds.

**Fig. 4:**
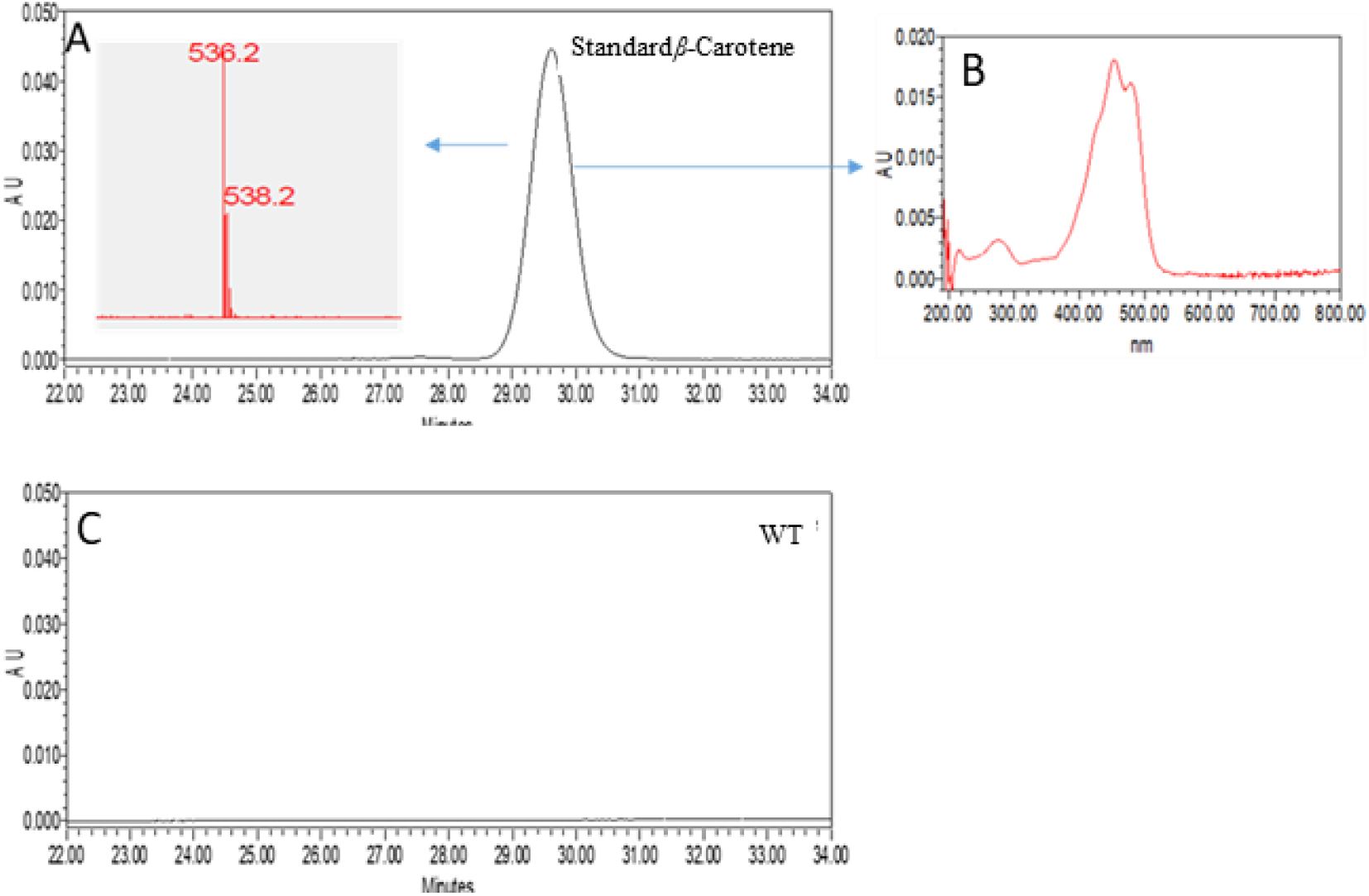
HPLC analyses of *β*-carotene from rice seeds. (A) reference *β*-carotene; (C) no *β*-carotene detected in wild type seeds

**Fig. 5:**
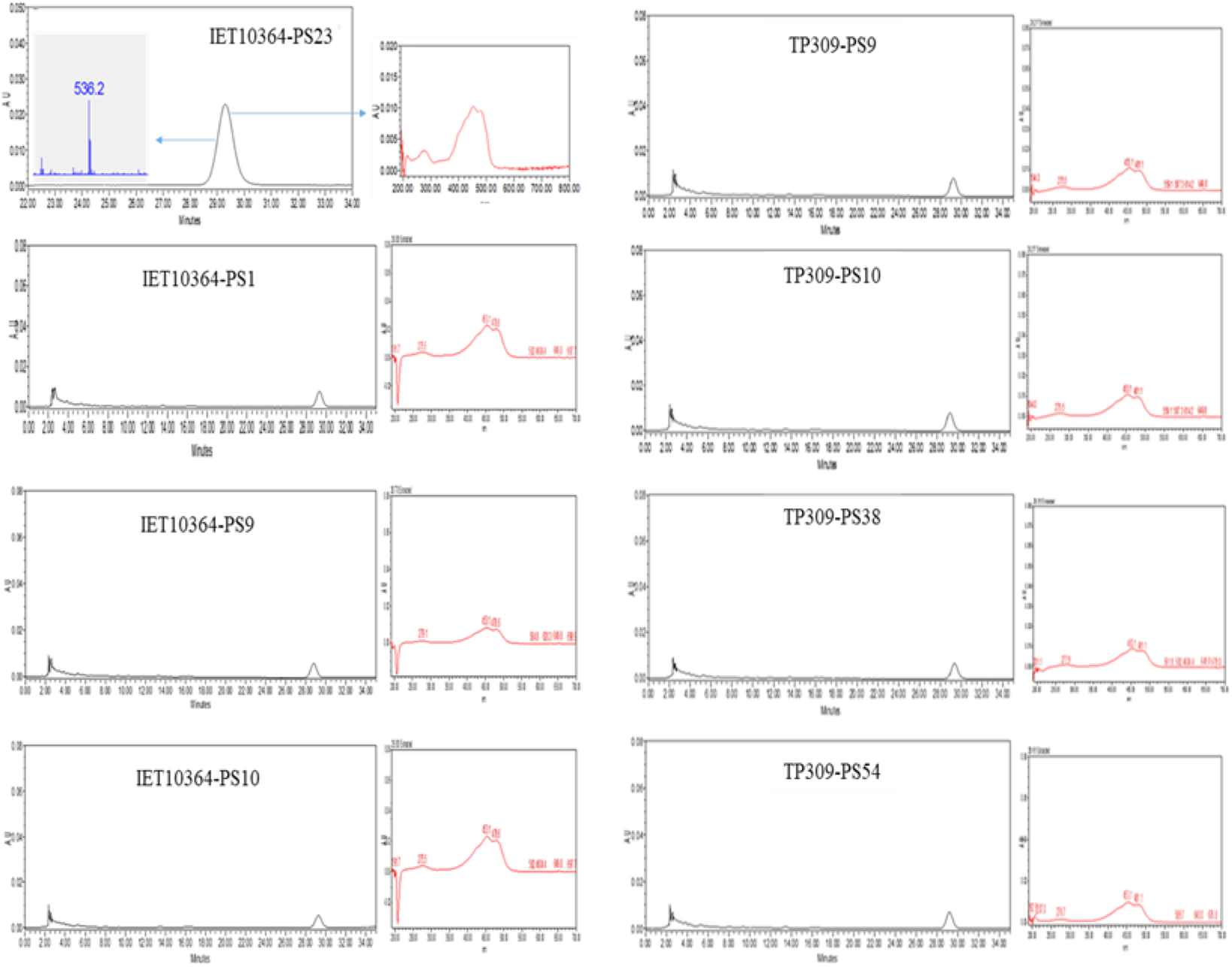
Chromatogram showing the presence of β-carotene content in transgenic seeds of IET10364 and TP309 lines seeds.

**Table 1.** Amount of β-carotene in wild type and transgenic rice lines.

| Plant Sample | β-carotene concentration ($\mu\text{g/g}$) |
| --- | --- |
| IET10364 WT | Not detected |
| IET10364-PS4 | 4.87 $\mu\text{g/g}$ |
| IET10364-PS23 | 13.54 $\mu\text{g/g}$ |
| TP309 WT | Not detected |
| TP309-PS55 | 2.82 $\mu\text{g/g}$ |
| TP309-PS7 | 8.20 $\mu\text{g/g}$ |

### Generation and verification of wheat transgenic lines

We developed 16 transgenic lines in wheat cv. CPAN1676 using *A. tumefaciens* strain EHA105 carrying binary vector *pANIC6B:Glu1D1-TaOr^His110^.* GUS histochemical assay was performed during callus formation to check if the transformation is successful. The putative transgenic T0 plants were removed from the non-transgenic plant using HPT, gene specific PCR (Supplementary Fig. S3C) and GUS histochemical assay (Supplementary Fig. S2B).

### qRT-PCR expression analysis of T2 developing seeds

Similar to rice, qPCR analysis was performed to analyze the expression pattern of transgene in T2 developing seeds of CPAN1676 at 20DAA stage. A similar relative expression of *TaOr^His110^* gene was found in various transgenic lines of CPAN1676 as compared to WT control (Fig 7).

**Fig. 6:**
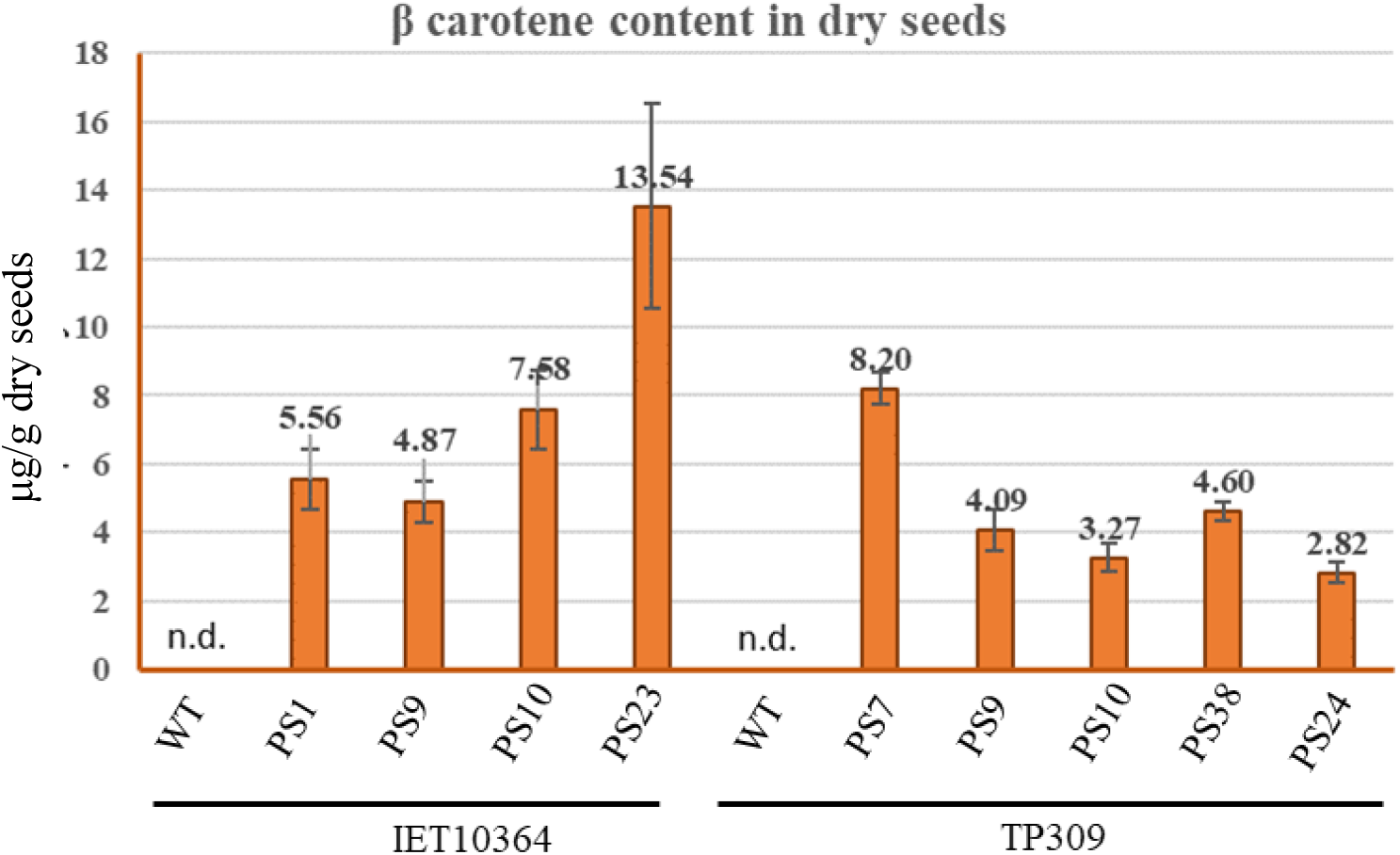
β-carotene concentration in IET10364-*Glu1D1TaOr^His110^* and TP309-*Glu1D1TaOr^His110^* (T1) transgenic grains and respective wild type control. On X-axis-WT and transgenic lines, Y axis-average value of β-carotene in µg/g seeds. Error bars denote the standard deviation

**Fig. 7:**
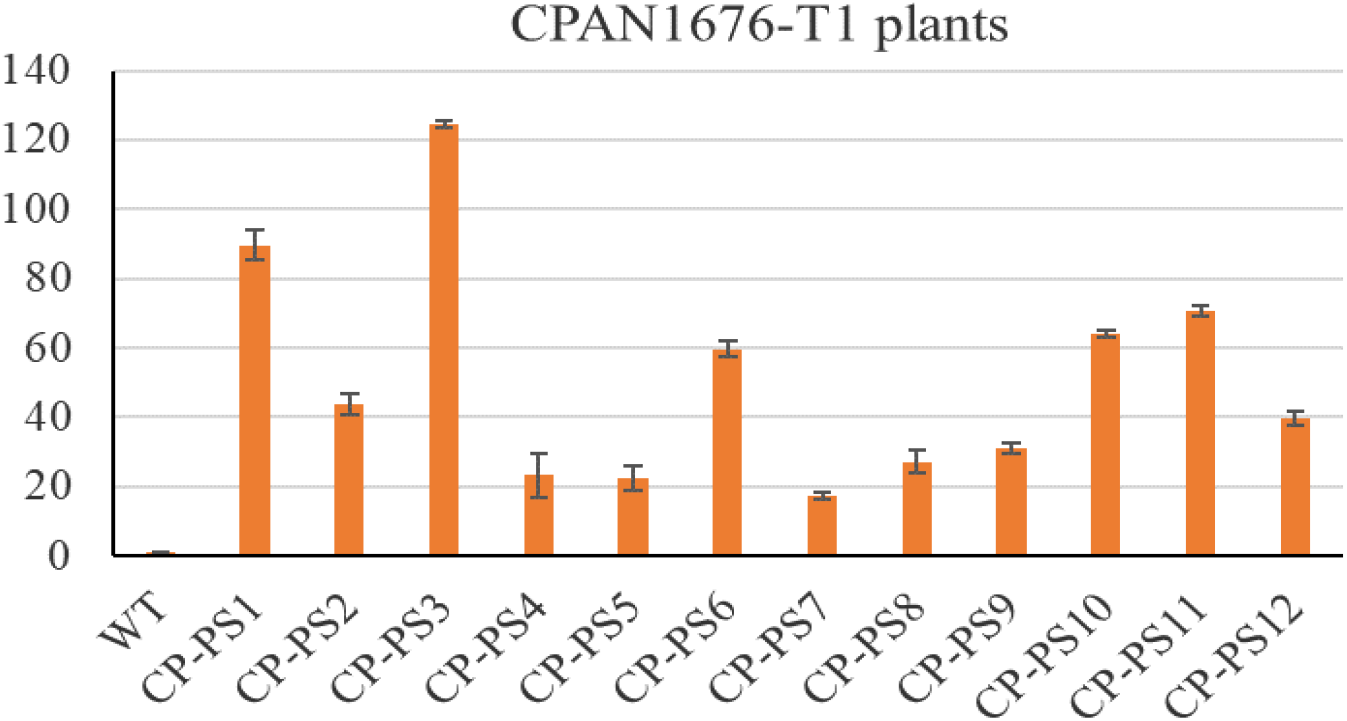
Relative expression of *TaOR* gene in CPAN1676-*Glu1D1TaOr^His110^* transgenic developing seeds (T1) with WT control. WT (CPAN1676 Wild-type control), CP-PS1 to CP-PS12are transgenic (T1) seeds. On X-axis-WT and transgenic wheat seeds, Y-axis-relative expression of gene. Error bars denote the standard deviation.

### Estimation of β-carotene concentration in CPAN1676-*Glu1D1TaOr^His110^* (T2) wheat grains

Estimation of β-carotene concentration of three lines each of CPAN1676-*Glu1D1TaOr^His110^* and flour from mix seeds of transgenic (T2) wheat grains was performed through HPLC. A clear distinction in color between powdered and intact seeds of transgenic and wild type seeds can be observed (Fig. 10A, B). The increase in β-carotene concentration of transgenic grains of CPAN1676 ranged from 3.67 µg/g (CP-PS3) to 7.18 µg/g (CP-PS11) as compared to 0.78 µg/g of WT control (Fig. 11A), which is approximately 10-fold. Similarly, approximately 7-fold increase in β-carotene concentration was found in powdered pooled transgenic grains from different lines of CPAN1676 as compared to powder from WT CPAN1676 grains (Fig.8)

**Fig. 8:**
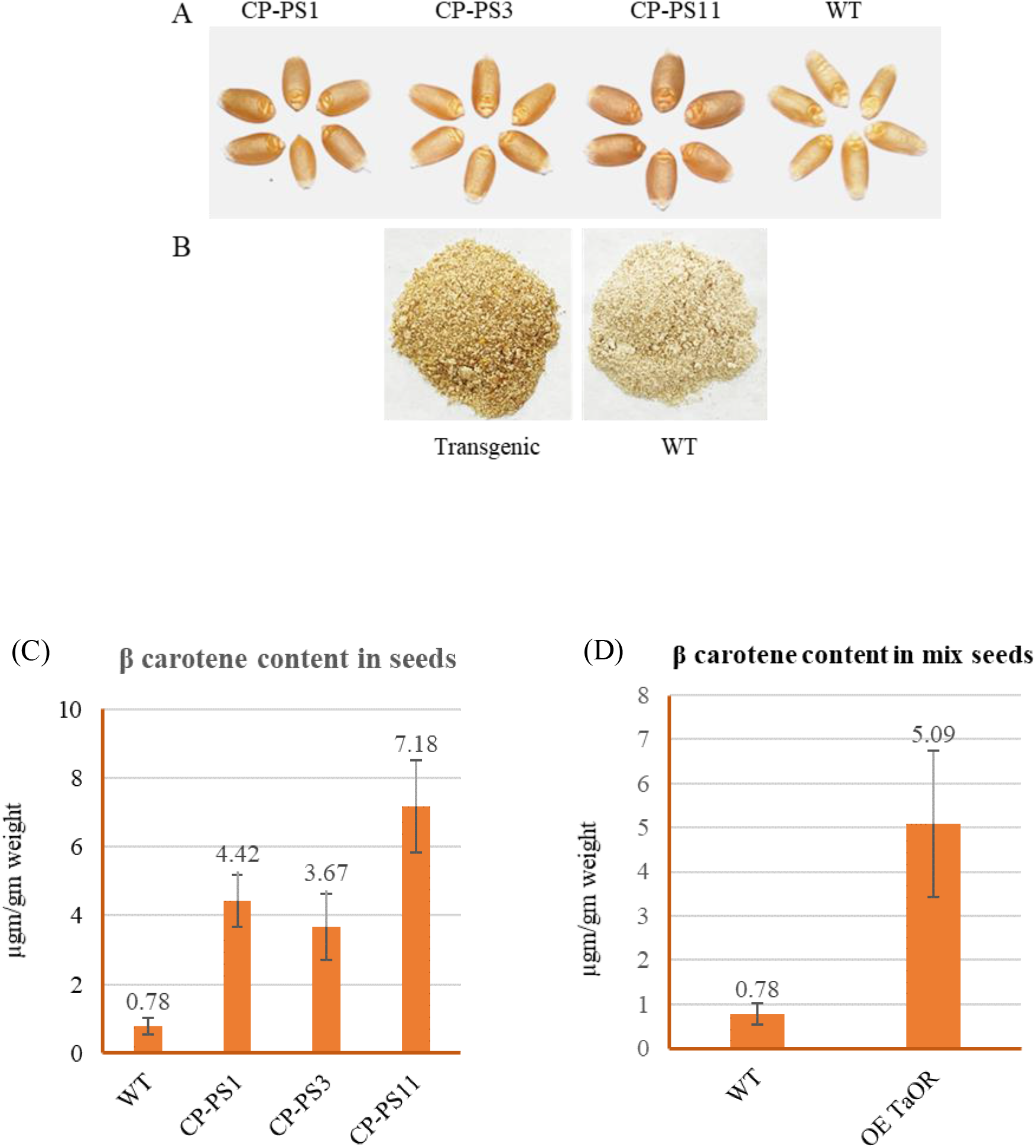
(A)Comparison of transgenic seeds of different lines of CPAN1676-*GluTaOr^His110^* with wild type seeds of CPAN1676. (B) Difference in the flour of transgenic and WT seeds. Estimation of β-carotene (C) β-carotene content in CPAN1676-*Glu1D1TaOr^His110^* (T1) transgenic grains with wild type control of CPAN1676. (D)β-carotene content in pooled grains of CPAN1676-*Glu1D1TaOr^His110^* (T1) transgenic grains with wild type control of CPAN1676. On X-axis-WT and transgenic lines, Y axis-average value of β-carotene in µg/g. Error bars denote the standard deviation

### *TaOr^His110^* overexpression significantly altered the expression of carotenoid metabolism-related genes in T1 seeds of IET10364-*Glu1D1TaOr^His110^*

To determine whether the accumulation of β-carotene in rice grains of *TaOr^His110^* overexpressing plants is correlated with altered expression of carotenogenesis-related genes, we analyzed the transcript levels of carotenoid biosynthesis genes in WT and *TaOR^His110^* plants by qRT-PCR. The results showed that the phytoene synthase (*PSY*), phytoene desaturase (*PDS*), zeta-carotene desaturase (*ZDS*), lycopene β-cyclase (*LCY-β*), carotenoid isomerase (*CRTISO*), β-carotene hydroxylase (*β-OH*), violaxanthin de-epoxidase (*VDE*), and zeaxanthin epoxidase (*ZE*) genes were expressed to higher levels in *TaOr^His110^*transgenic grains of IET10364 and TP309 than in their respective WT grains. The expression pattern of carotenoid biosynthesis genes is different in all the lines, but in the case of IET10364-PS23 plant, all genes showed significant upregulation as compared to WT, which is in accordance with more β-carotene accumulation in this line. Similarly, in case of TP309 transgenic grains, TP-PS7, TP-PS38 and TP-PS55 showed significant upregulation in expression of carotenoid biosynthesis genes (Fig 9).

**Fig. 9:**
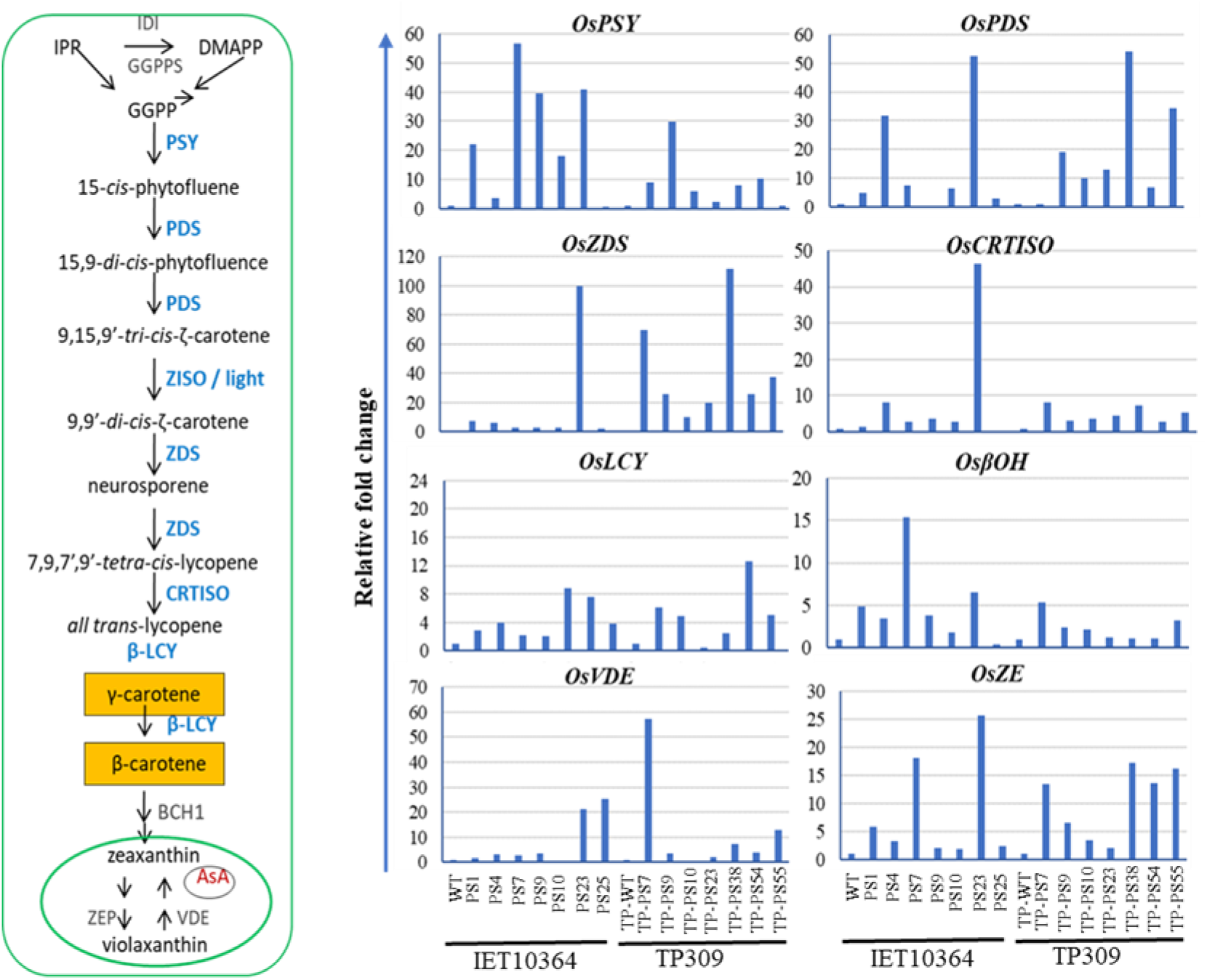
Expression patterns of various carotenoid biosynthesis genes. Expression patterns of various genes involved in carotenoid biosynthesis in grains of WT and *TaOr^His110^*-overexpressing plants. X-axis shows specified transgenic lines, Y-axis shows relative fold change in expression compared to respective WT. *PSY*, phytoene synthase; *PDS*, phytoene desaturase; *ZDS*, ζ-carotene desaturase; *LCY-β*, lycopene β-cyclase; *CRTISO*, carotenoid isomerase; *β-OH*, β-carotene hydroxylase; *VDE*, violaxanthin de-epoxidase; *ZE*, zeaxanthin epoxidase

## Discussion

To maintain metabolism, all living species require necessary micronutrients, which humans acquire through their diet (Stefanache et al.2023). However, basic cereals like wheat and rice have inadequate levels of micronutrients, particularly iron (Fe) and vitamin A, moreover milling removes most of their content, leaving nearly nothing. Furthermore, micronutrient deficiencies occur in locations where the human diet consists primarily of cereals. Since almost one third of our population lives below poverty level, providing a balanced diet is a herculean task. In view of the above facts, it is particularly important to improve the nutritional quality of rice and wheat by increasing its provitamin A content through conventional and/or genetic engineering approaches.

Increased provitamin A accumulation has been reported to be caused by a novel *Or* gene isolated from a high β-carotene orange cauliflower mutant (Lu et al, 2006). *Or* encodes a DnaJ Cys-rich zinc finger domain containing protein, which is highly conserved among divergent plant species (Lu et al., 2006). It has been already shown that an *Or* gene mutant with insertion of a copia-like LTR retrotransponson or alteration of a single amino acid in a wild-type *OR* greatly enhanced its ability to promote carotenoid accumulation in normally low-pigmented tissues (Zhou et al., 2008; Yuan et al., 2015). Yuan et al. (2015) demonstrated that alteration of a single amino acid (*Or*^His^ and *Or*^Ala^) in a wild-type *OR* by site-directed mutagenesis promoted high levels of carotenoid accumulation which further suggests its potential as new genetic tool for crop carotenoid enhancement. Lopez et al. (2008) demonstrated that manipulating sink formation offers an alternative technique for increasing carotenoid content in food crops by the successful application of the *Or* gene to raise carotenoid levels in transgenic potato tubers. However, the functional characterization of this *Or* gene in wheat and rice have not been explored so far. Therefore, in the present study, we aimed to assess the *in vivo* effects of *TaOr* gene in increasing β-carotene content in rice and wheat grains through endosperm specific expression.

Plant Or proteins contain an N-terminal region of unknown function, transmembrane domains, and a C-terminal DnaJ-like domain which are highly conserved among plant species (Kim et al., 2019; Lu et al., 2006), suggesting that they play an important role in plant growth and development. However, Or proteins have been functionally characterized in only a few plant species, including cauliflower, melon, sorghum, *Arabidopsis*, and sweet potato (Lu et al., 2006; Tzuri et al., 2015; Yuan et al., 2015; Bai et al., 2014; Kim et al., 2013). The cauliflower Or protein localizes both in plastids and nuclei (Lu et al., 2006; Zhou et al., 2015). Interestingly, it was reported in sweet potato that Or protein translocate from nuclei to chloroplasts in response to heat stress (Park et al., 2016). In *Arabidopsis,* Or protein localization has been reported either as both in chloroplasts and nucleus (Sun et al. 2019; Zhou et al. 2015) or in chloroplasts only (Shan et al., 2022). In this present study, TaOr protein overexpressed in tobacco was detected only in the chloroplasts of leaf epidermal cells (Fig. 1B), suggesting that Or is mainly localized in the chloroplast in plant leaves, which induce carotenoid synthesis. The TaOr protein contains two trans-membrane domains and a DnaJ cysteine-rich zinc finger domain which is similar to the Or proteins in other plant species. DnaJ proteins belong to a large protein family that is characterized by different subcellular localizations (Miernyk, 2001). However, many DnaJ proteins localize to chloroplasts, for example, in *Arabidopsis* 19 DnaJ proteins are reported to be localized in chloroplast (Chiu et al., 2013). The DnaJ proteins are heat shock proteins that are important for chloroplast growth, photosynthesis, and abiotic stress tolerance, implying that Or gene expression might increase plants’ ability to adapt to environmental challenges (Albrecht et al., 2008; Wang et al., 2014, 2015).

*TaOr* was cloned from wheat and site-directed mutagenesis was done to change the amino acid Arginine at 110^th^ position to Histidine. This *TaOr^His110^* was cloned under the seed-specific promoter *Glu-1D1* to obtain its expression in endosperm of rice and wheat grains. Overexpression of *TaOr^His110^*in both cultivars of rice and one cultivar of wheat changed the color of endosperm from white to light orange/ brown (Fig 3 and 8). These results suggest that overexpression of *TaOr^His110^*under a seed-specific promoter increased the carotenoids contents, especially β-carotene, in endosperm of both rice and wheat. In comparison with the leaf tissue, higher accumulation of β-carotene in the endosperm of *TaOr^His110^* expressing lines supports the suggestion that a tissue-specific promoter would be a better strategy for carotenoid metabolism engineering in rice and wheat, not only for its spatio-temporal specificity of expression but also for its strong promoter activity. In this study, we showed that *TaOr^His110^* overexpression in wheat and rice endosperm increased the accumulation of β-carotene.

Previously, overexpression of the golden SNP-carrying *CmOr* mutant resulted in increased carotenoid content and orange-colored melon fruit phenotypes as compared to green-colored melon in wild type (Tzuri et al., 2015). By overexpressing the *IbOr* gene containing the same mutation, (Kim et al., 2019) demonstrated a rise in β-carotene content and improved abiotic stress tolerance in sweet potato. A significant increase in potato was observed by expression of sweet potato and cauliflower *Or* genes (Lopez et al., 2008; Li et al., 2012; Cho et al., 2016).

Or stimulates chromoplast formation in plants which provides metabolic sink for carotenoid accumulation (Cazzonelli and Pogson, 2010; Li et al., 2012). Overexpression of *Arabidopsis Or* gene increased carotenoid concentration in transgenic rice and corn by promoting chromoplast formation (Bai et al., 2016; Berman et al., 2017).

Further, β-carotene concentration was measured in transgenic rice and wheat grains with WT control using HPLC. The β-carotene concentration of transgenic TP309-*Glu1D1TaOR^His110^* grains increased and ranged from 2.82 to 8.20 µg/g. The β-carotene concentration of IET10364-*Glu1D1TaOR^His110^* transgenic grains also increased and ranged from 4.87 to 13.54 µg/g and in accordance with reports of transgenic rice (Paine et al., 2005; Ye et al., 2000). In WT seeds of TP309 and IET10364, the amount of β-carotene was below the detection limit. Previously, Ye et al. (2000) reported invention of Golden rice 1 containing approx. 1.6 µg/g β-carotene by overexpression of *psy* (*phytoene synthase*) gene from daffodil and two other genes *crtI* (*phytoene desaturase*) and lcy (*lycopene cyclase*) from *Erwinia uredovora*. Subsequently, Golden rice 2 containing 37 µg/g β-carotene was produced by Paine et al. (2005) by replacing daffodil *psy* by its homolog from maize. Tian et al. (2019) utilized metabolic engineering to enhance carotenoid biosynthesis in rice endosperm by improved flux through the mevalonate route, due to expression of *tHMG1*, *Zea maize PSY1*, and *PaCRTI* genes. Similarly, increase in β-carotene concentration of CPAN1676-*Glu1D1TaOR^His110^* transgenic wheat grains ranged from 3.67 to 7.18 µg/g as compared to 0.78 µg/g in WT grains which is approximately 10 folds higher than WT. Cong et al. (2009) reported 10.8 folds higher β-carotene levels in transgenic wheat as compared to wild-type, by the transformation of the maize *PSY*1 and *Crt*I genes. Similarly, Wang et al. (2014) co-expressed the bacterial *Crt*B and *Crt*I genes which resulted in 8-fold increase in β-carotene level of wheat.

We investigated the expression of endogenous carotenogenic genes to learn more about the link between increased carotene accumulation in transgenic plants and up-regulation of endogenous carotenoid biosynthesis genes, including phytoene synthase (*PSY*), phytoene desaturase (*PDS*), zeta-carotene desaturase (*ZDS*), lycopene β-cyclase (*LCY-β*), carotenoid isomerase (*CRTISO*), β-carotene hydroxylase (*β-OH*), violaxanthin de-epoxidase (*VDE*), zeaxanthin epoxidase (*ZE*) genes, by RT-qPCR upon their putative regulation as a result of transgene expression. In transgenic TP309 seeds, the transcripts of OsPDS, OsLYCE, OsZDS, OsVDE and OsZE were increased to significantly higher levels of upto 110-fold relative to their expression in WT plants (Fig. 9). Similarly, in transgenic IET10364 seeds, the expression of almost all the biosynthetic genes was increased (Fig. 9), correlating with an increased β-carotene content. The results are similar to the report provided by Tian et al. (2019) which suggests upregulation of carotenoid biosynthetic genes leads to increase in carotenoid content. Welsch et al. (2018) demonstrated the formation of membrane-complex PSY-Orange, which is responsible for carotenogenesis and degradation of non-associated and misfolded PSY by Clp proteases. As a result, Orange protein is an important enzyme in the regulation of carotenoid production (Chayut et al., 2017; Welsch et al., 2018)

Keeping this in mind, we identified the potential of the *TaOr* gene which is a DnaJ like molecular chaperone with cysteine rich residues in rice and wheat by developing overexpressing transgenic lines under seed-specific *Glu1-D1* promoter from *Triticum aestivum*. Our study found significant increase in β-carotene concentration in both TP309 (*japonica*) and IET10364 (*indica*) variety of transgenic rice along with CPAN1676 variety of wheat. Future studies can be performed to identify the interacting partners of *Or* gene through chromatin immunoprecipitation assays in overexpressing transgenic lines. *TaOr* gene can further be analyzed along with co-expression of important carotenoid biosynthesis genes for increase in total carotenoid content. Also, a combination with other seed-specific promoter can be checked for increase in β-carotene content in endosperm. In addition, the analysis of the transgenic lines obtained in this study can be performed for important abiotic stresses such as heat, salinity and drought stresses.

